# Extinction cascades, community collapse, and recovery across a Mesozoic hyperthermal event

**DOI:** 10.1101/2022.06.13.495894

**Authors:** Alexander M. Dunhill, Karolina Zarzyczny, Jack O. Shaw, Jed W. Atkinson, Crispin T.S. Little, Andrew P. Beckerman

## Abstract

Biotic interactions and community structure are seldom examined in mass extinction studies but must be considered if we are to truly understand extinction and recovery dynamics at the ecosystem scale. Here, we model shallow marine food web structure across the Toarcian extinction event in the Cleveland Basin, UK using a trait-based inferential modelling framework. First, we subjected our pre-extinction community to extinction cascade simulations in order to identify the nature of extinction selectivity and dynamics. Second, we tracked the pattern and duration of the recovery of ecosystem structure and function following the extinction event. In agreement with postulated scenarios, we found that primary extinctions targeted towards infaunal and epifaunal benthic guilds reproduced the empirical post-extinction community. These results are consistent with geochemical and lithological evidence of an anoxia/dysoxia kill mechanism for this extinction event. Structural and functional metrics show that the extinction event caused a switch from a diverse, stable community with high levels of functional redundancy to a less diverse, more densely connected, and less stable community of generalists. Ecological recovery appears to have lagged behind the recovery of biodiversity, with most metrics only beginning to return to pre-extinction levels ∼7 million years after the extinction event. This protracted pattern supports the theory of delayed benthic ecosystem recovery following mass extinctions even in the face of seemingly recovering taxonomic diversity.

---

Earth has experienced a number of mass extinction events that have shaped evolutionary history, not only by the dramatic loss of species over relatively short periods of time but also by repeatedly restructuring ecosystems^1^. Many mass extinctions have been linked to rapid releases of greenhouse gases into the atmosphere by large igneous province (LIP) volcanism, which led to a cascade of environmental effects, including rapid global warming, ocean anoxia, and ocean acidification^2^. Thus, palaeobiologists have long viewed mass extinctions as the quintessential example of a Court Jester driver of macroevolution, whereby external abiotic environmental pressures act as the predominant driver of macroevolution across long geological timescales^3^. However, community ecological theory suggests that biotic interactions, and thus Red Queen processes^4^, would have also played a major role during mass extinction events. Many victims of past extinction events were unlikely to have gone extinct as a direct effect of abiotic stress, but probably did so in response to cascading secondary effects via the loss of key prey sources^5^.

Macroecological studies of selectivity^6,7^, functional diversity loss^8,9^, and recovery^10,11^ indicate that warming-related mass extinctions are associated with of latitudinal extinction selectivity^6,12,13^ as well as preferential loss of taxa vulnerable to hypercapnia, anoxia and acidification, and photosymbiotic taxa^6,13-15^. All of these patterns can be qualitatively linked to one of the aforementioned abiotic effects of LIP eruptions, but it remains incredibly difficult to ascertain the full range of, or relative impact of, abiotic stressors that contributed to a particular mass extinction event.

Ecological theory suggests that extinction dynamics are most effectively studied within a community framework where details about interactions among taxa may accentuate or buffer the responses of individual taxa from direct (i.e. primary) and cascading ‘secondary’ extinctions^16^. In fact, many victims of mass extinctions are unlikely to have become extinct as a direct effect of abiotic stress, but probably did so in response to cascading secondary effects^5^. Biotic interactions are seldom considered in mass extinction studies (but see^17-19^) as it is very difficult to ascertain consumer-resource and other interactions between long extinct organisms. However, if we are to truly understand mass extinction dynamics, we must quantify such interactions as many prominent extinction patterns, such as the high levels of extinction amongst pelagic predators^6^, are difficult to explain in the absence of extinction cascades through communities^20^.

In addition to the study of mass extinction causality, magnitude, and selectivity, extinction recovery has also become an intensely studied topic over the past decade^21^, driven in part by the desire to understand how long the planet may take to recovery from the current “Sixth Mass Extinction”^22^. Studies based on taxonomic and functional diversity suggest full ecosystem recovery can take anywhere between 0.7 to 50 million years from the largest mass extinctions^10,11,23^, but it is possible that ecosystem function could recover despite persistent low levels of biodiversity. Thus, studies of extinction recovery could be greatly improved by adopting a community ecology approach that integrates across biodiversity, community structure and ecosystem function.

Here, we utilise a community ecology food web approach to model primary and secondary extinction dynamics and community recovery across the early Toarican extinction event (ETE; ∼183 Ma) from the Cleveland Basin, Yorkshire, UK. Specifically, we use ecological trait data to reconstruct plausible food webs. We then subject these food webs to several primary extinction scenarios that link event characteristics (e.g. dysoxia, acidification, warming) to traits. We use well established ecological modelling tools to evaluate patterns of secondary extinction, ultimately identifying several target traits and species whose sensitivity to the ETE event led to the ensuing post event community structure. Finally, we also look at empirical patterns of recovery from this extinction event, detailing changes in biodiversity, functional groups and community structure.

The ETE, is traditionally referred to as a second order extinction event^24^ (i.e. an extinction event that caused less than 40% generic extinction^25^) and resulted in the loss of around 26% of marine genera globally^26^. It is linked to the eruption of the Karoo-Ferrar Large Igneous Province^27^ which resulted in a globally distributed negative carbon isotope shift^28,29^, hyperthermal warming of up to 13ºC in the midlatitudes^30,31^, prolonged regional ocean dysoxia and anoxia^26,32-34^, and ocean acidification^35^.

In the Cleveland Basin and in much of the NW Tethyan basins of NW Europe the ETE is coincident with the deposition of finely laminated, organic-rich, black shales which signify persistent dysoxia/anoxia, at shallow depths on the continental shelf, termed the Toarcian Oceanic Anoxic Event (TOAE or Jenkins Event)^33,36^. The ETE resulted in the loss of around 60% of marine species within the Cleveland Basin (87% benthic species extinction) ^24,37^, with post-extinction benthic communities made up of low diversity/high abundance assemblages of hypoxia-tolerant and opportunistic species^26,34,38,39^. Recovery seems to occur in two pulses and, in total, takes as long as 7 million years in terms of both taxonomic and functional diversity^37,40^.

We use a data set of 38,670 occurrences of 162 species of marine invertebrates, fish, and trace fossils derived from years of detailed field studies^24,26,41,42^ of one of the most expanded Pliensbachian to Toarcian sections in the world to produce a series of community trophic networks (i.e. food webs) (Fig. 1). We aim to ascertain: (i) the most plausible set of traits and species impacted by primary extinction events and the nature of any secondary extinction cascades, that led to the post extinction community structure and diversity; (ii) the likely environmental trigger of the extinction cascades; and (iii) the pattern and duration of ecological recovery in the aftermath of the extinction event.

**Figure 1.**
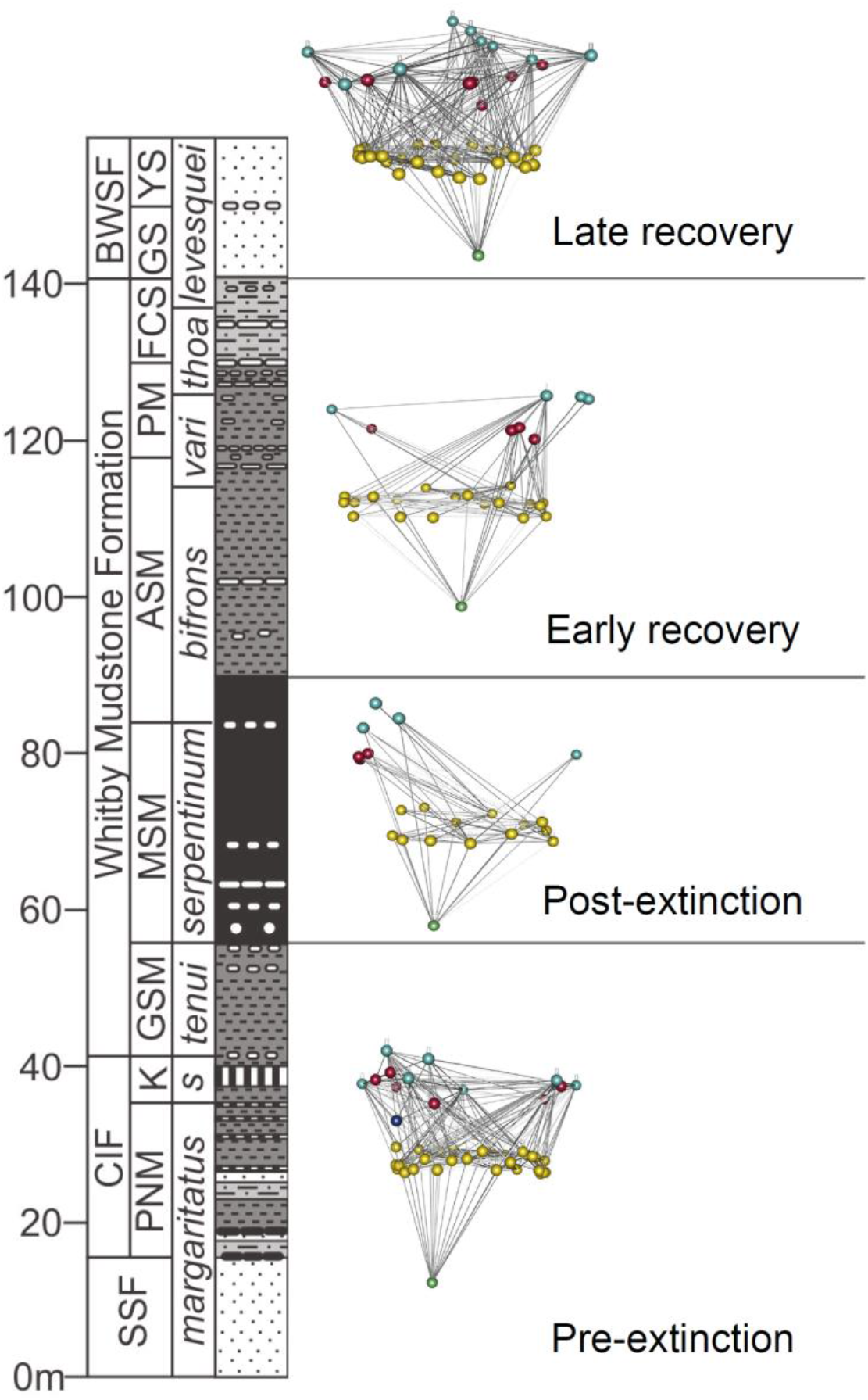
Stratigraphic column of the Pliensbachian-Toarcian (Lower Jurassic) of the Cleveland Basin at Ravenscar (North Yorkshire, UK) showing community food webs for pre-extinction, post-extinction, early recovery and late recovery intervals. Lithostrat column abbreviations need explanation. SSF = Staithes Sandstone Fomration; CIF = Cleveland Ironstone Formation; BWSF = Blea Wyke Sandstone Formation; PNM = Penny Nab Member; K = Kettleness Member; GSM = Grey Shales Member; MSM = Mulgrave Shales Member; ASM = Alum Shales Member; PM = Peak Mudstone Member; Fox Cliff Sandstone Member = ; GS = Grey Sandstone Member; YS = Yellow Sandstone Member; *s* = *spinatum*; *vari* = *variabilis*; *thoa* = *thouarsense*.

## Results and discussion

### The Early Toarcian extinction event

#### Empirical data: pre-extinction

The pre-extinction community is characterised by a diverse assemblage of benthic and pelagic taxa belonging to 48 different trophic guilds (Fig. 1; Fig 2A; see supplementary data for network metrics values for all communities). The community consists of a range of primary consumers (including suspension feeders, deposit feeders, miners, and grazers), intermediate predators (cephalopods, crustaceans, fish, gastropods, scaphopods), and top predators (larger cephalopods). The pre-extinction community is characterised by values of many common network metrics that are well within the bounds for typical modern day marine communities (e.g. connectance, generality, vulnerability, mean/max. trophic level etc.)^5,39,43^ (Fig 2B).

**Figure 2.**
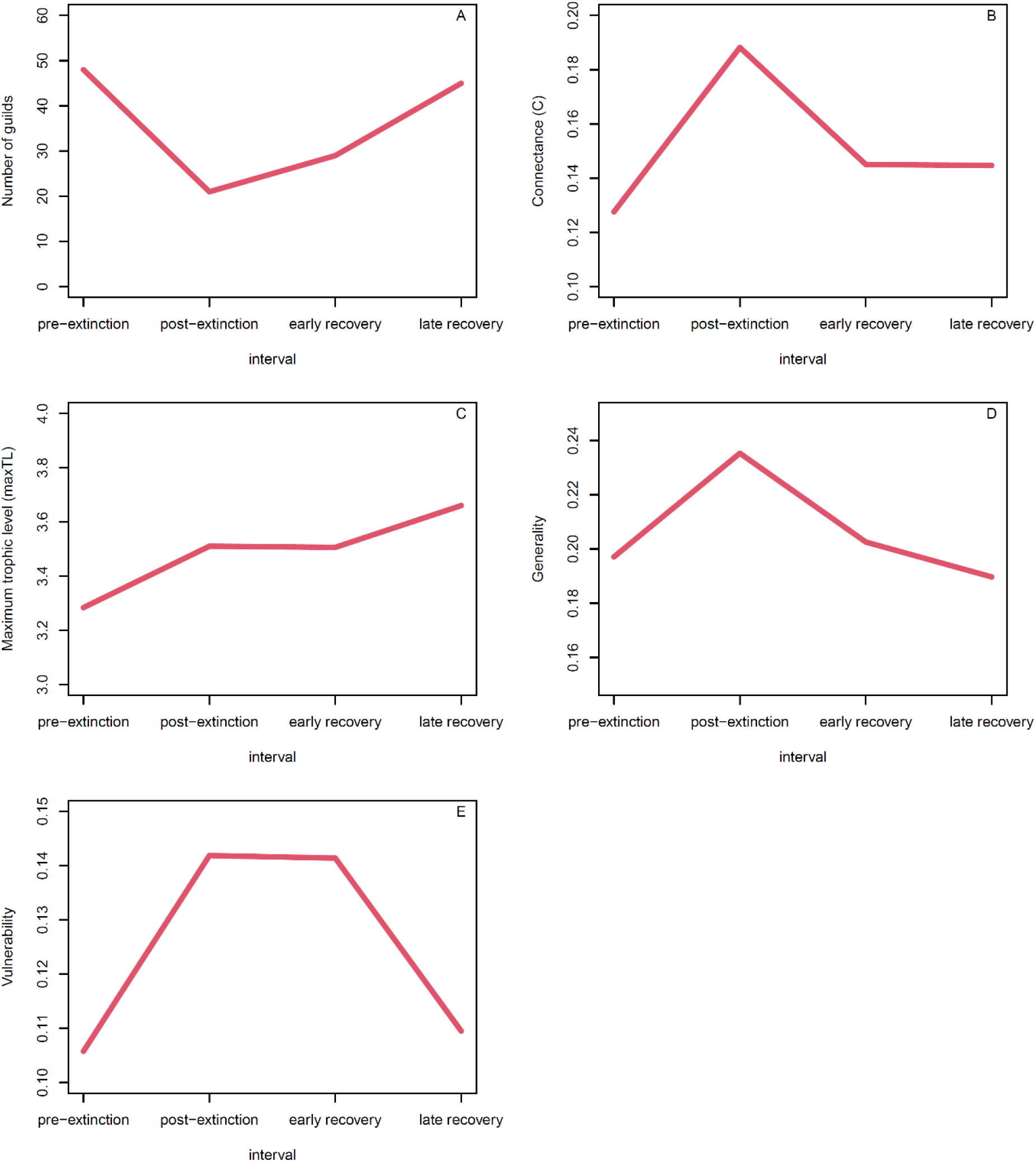
Structural food web metrics across the four intervals of the ETE; (A) guild richness; (B) connectance (C); (C) maximum trophic level (MaxTL); (D) generality; (E) Vulnerability.

#### Empirical data: post-extinction

The ETE causes a reduction in richness from 48 to 21 trophic guilds (Fig. 1; Fig 2A) and drove disproportionate losses amongst infaunal, large, highly motile, and predatory benthic guilds (∼80% benthic species extinction^24,37^). Post-extinction benthic assemblages were dominated by low-diversity/high-abundance communities of very small, epifaunal, suspension-feeding bivalves, most notably the presumably low-oxygen-tolerant opportunistic species *Bositra buchii* and *Pseudomytiloides dubius*^34^. Although previous research has suggested that pelagic taxa we also affected by the ETE^44^ (including high mortality amongst pelagic faunas and cephalopods migrating away from the Cleveland Basin in response to high temperatures and low food supply), at the structural level, pelagic elements of the community appear much less affected.

The severe benthic losses at the guild level are in contrast to global studies of mass extinctions which have postulated that, although species losses are typically severe, mass extinctions rarely cause global guild-level extinction^8,45^. However, the loss of >50% of guilds in this local, mid-latitude setting is in keeping with regional studies of extinction across the ETE^24,26,37,46-49^, that have shown species and guild-level losses to be much higher in the mid latitudes (i.e. NW Tethys and NE Panthalassan margin) than globally^6,45^. The post-extinction benthic assemblage is dominated by a low-diversity/high-abundance community of small-bodied, epifaunal, suspension feeders with apparent selectivity against larger benthic taxa with more active modes of life. Together with lithological and geochemical evidence, this strongly supports the theory of a dysoxia/anoxia driven extinction in the Cleveland Basin^41^.

Overall community connectivity (i.e. connectance) increased after the ETE (Fig. 2B) which also corresponds with an increase in generality, vulnerability, and maximum trophic level within the community (Fig. 2B-E). Together with a reduction in the number of linear chains within the food web (S1), the levels of omnivory (S2) and both apparent (S4) and direct competition (S5) (Fig. 3A-D) this suggests that the post-extinction community consists of fewer guilds that are more generalistic in their feeding habits and thus taxa are more closely linked to one another via consumer-resource interactions than taxa in the pre-extinction community. Selective extinction of benthic taxa, which are predominantly lower- and intermediate-level consumers, led to the food web becoming taller and thinner (i.e. fewer linear chains) with more restricted energy flows, fewer lower-level consumers, and increased predation pressure on the remaining lower trophic level taxa (i.e. increased vulnerability). This also led to reduced direct completion (S5) as benthic predators disappear from the community and reduced apparent competition (i.e. predator choice; S4) as the extinction wiped out the majority of the benthic guilds such that pelagic predators had fewer prey options.

**Figure 3.**
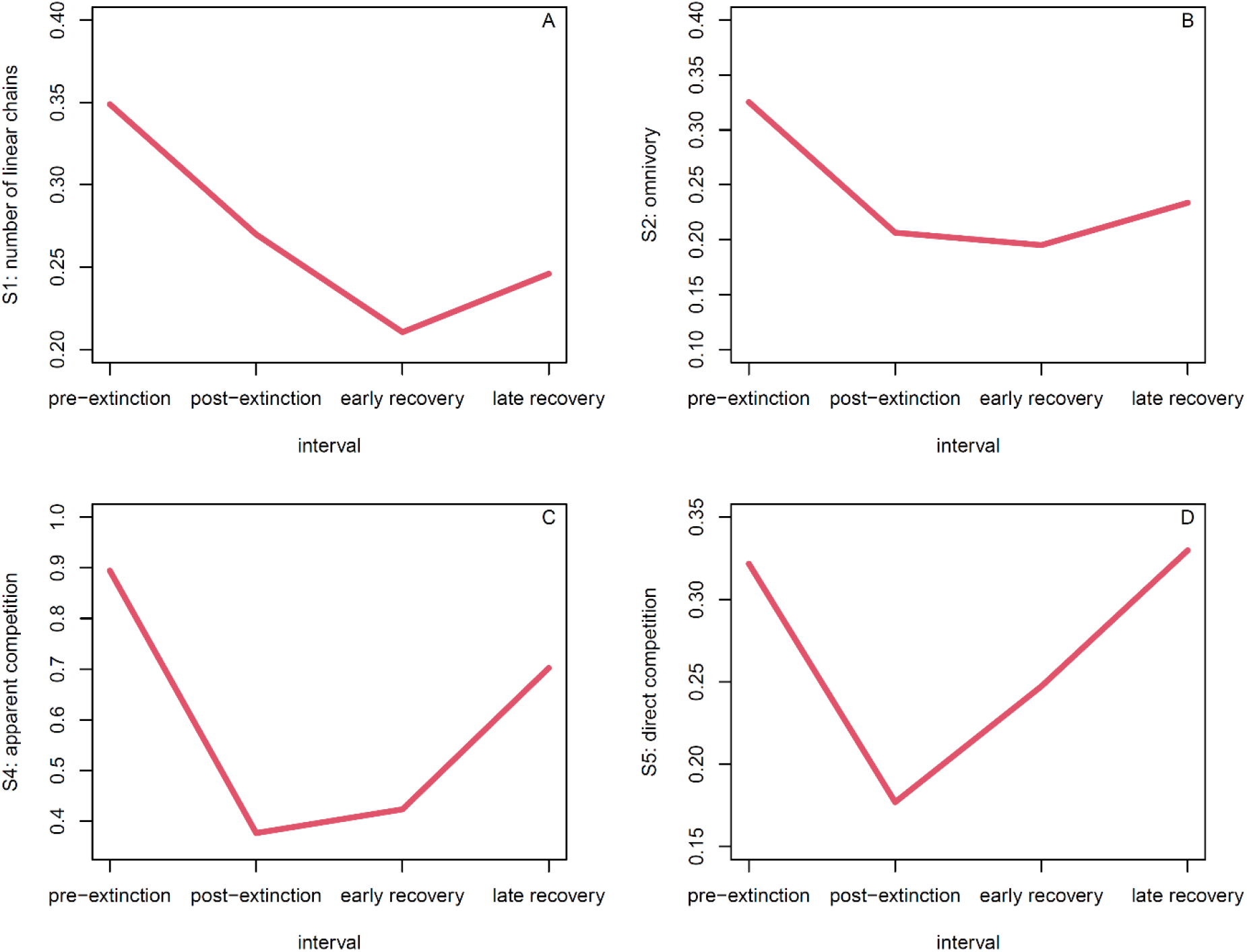
Functional food web motifs across the four intervals of the ETE; (A) S1: number of linear chains; (B) S2: number of omnivory motifs; S4: number of apparent competition motifs; (D) S5: number of direct competition motifs.

The post-extinction community also displays lower levels of omnivory as the intermediate consumers in the benthic realm go extinct, meaning that top predators are now feeding across fewer trophic levels. This pattern of densely connected, species-poor communities of opportunists/generalists is consistent with evidence from palaeoecological interpretations of the fossil record (i.e. low-diversity/high-abundance communities of opportunistic species)^34,37^ and other instable post-mass extinction food webs reconstructed from the fossil record^18,19,50^.

This post extinction community bears a broadly similar structure to that of modern low-diversity communities dominated by generalists^43^. Although there is some empirical evidence that higher connectance leads to higher community robustness (i.e. more stable communities that are less likely to collapse)^51,52^, the taller, thinner web with reduced omnivory values of the post-extinction web suggests instability as energy flows are contingent on very specific pathways and presents a rivet-hypothesis scenario where the removal of a few well-connected guilds could lead to wholesale ecosystem collapse^52,53^. The removal of intermediate consumers (i.e. benthic crustaceans, gastropods, scaphopods) increases the vulnerability of the few remaining species of primary consumers in the benthic realm as they continue to be predated by a similar number of pelagic higher level consumers (i.e. cephalopods) as seen in the pre-extinction community. This change in the tiering “pyramid” suggest community-level patterns reflect patterns seen in the global ecosystem in the aftermath of mass extinction events that have strong bottom-water dyoxia/anoxia drivers^11^ as well as communities in present-day deoxygenated areas^54^.

#### Extinction dynamics

We simulated, with replication, 13 different primary extinction scenarios (see Methods). Each of these scenarios generated unique signatures of primary and secondary extinction. All scenarios with identical starting guild richness (i.e. 48) were terminated when the simulations reached the post-extinction richness of 21. We calculated multiple structural metrics and a True Skill Statistic (TSS)^55^ to compare the simulated post-extinction events to the actual post-extinction event.

Three of the simulated extinction scenarios produced communities that were much more similar to the empirical post-extinction community than the random primary extinction scenario (Fig. 4A). According to the TSS scores, primary extinction selectivity based on tiering, with strongest extinction selectivity against infaunal taxa, gave by far the closest replication of the empirical post-extinction community (Fig. 4A). Targeting generality was the second most plausible scenario. The other 10 were largely indistinguishable from random (Fig 4a).

**Figure 4.**
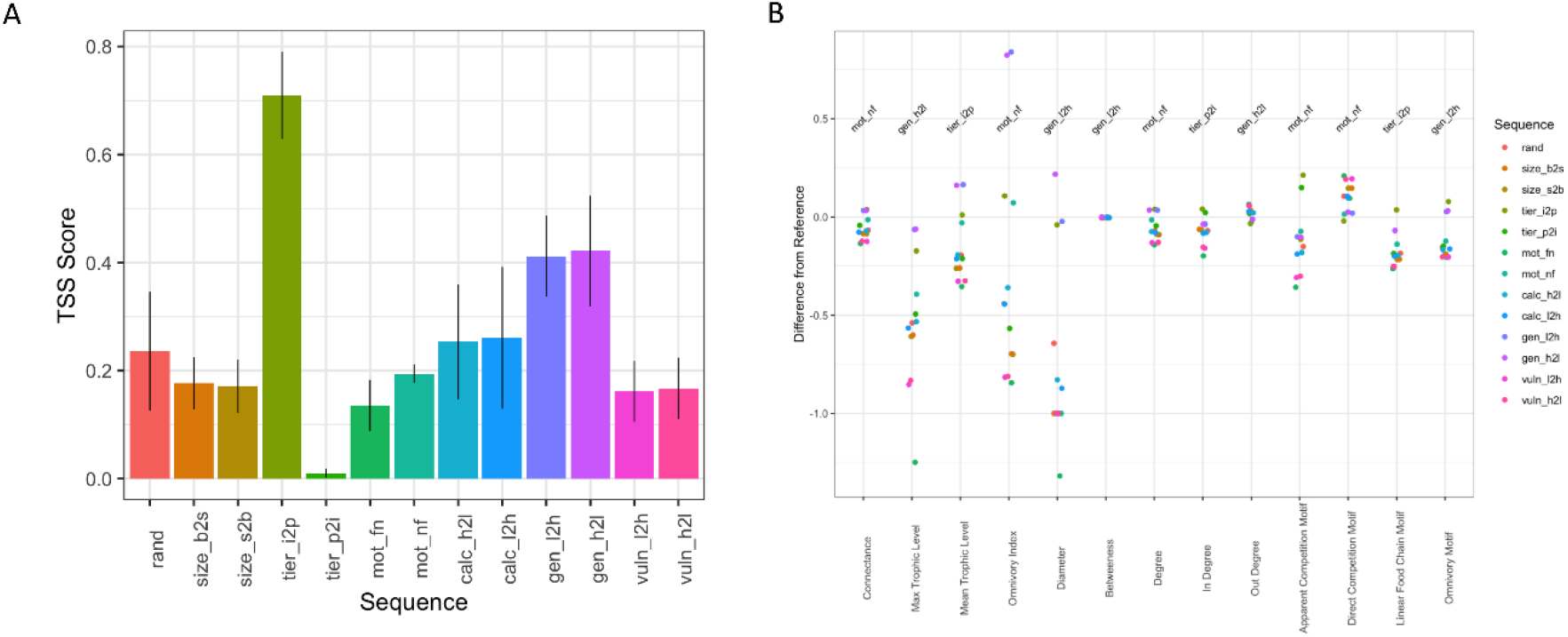
Results of secondary extinction cascade simulations showing similarity between simulated and empirical pre-extinction food web structure and function via; (A) True Skills Statistic showing the similarity in node presence/absence between simulated scenarios and empirical post-extinction community; (B) similarity in structural food web metrics and food web motifs between simulated scenarios and empirical post-extinction community (see Table 1 for sequence definitions).

#### Focus: tiering

The similarities between the empirical post-extinction community and the simulated extinction scenario based on primary extinction selectivity against infaunal and benthic taxa suggests that the strongest extinction selectivity was placed upon taxa living within the sediment or on the seabed in the benthic realm. This pattern matches (i) the empirical data where the majority of extinction occurs within the benthic realm (with almost all infaunal guilds disappearing) and (ii) an anoxia/dysoxia kill mechanism dictating that bottom waters would contain less oxygen than sea surface layers with the capacity for continuous gas exchange with the atmosphere. This result is in line the lithological evidence for anoxia/dysoxia (i.e. organic-rich black shales) within the Cleveland Basin^38^ and provides confidence that our method is performing well.

**Table 1.** Definitions of extinction cascade simulation sequences.

| Sequence | Definition |
| --- | --- |
| rand | Primary extinctions targeted in a random fashion |
| size_b2s | Primary extinctions targeted at larger guilds |
| size_s2b | Primary extinctions targeted at smaller guilds |
| tier_i2p | Primary extinctions targeted at infaunal > epifaunal > pelagic guilds |
| tier_p2i | Primary extinctions targeted at pelagic > epifaunal > infaunal guilds |
| mot_fn | Primary extinction targeted at most motile guilds |
| mot_nf | Primary extinction targeted at least motile guilds |
| calc_h2l | Primary extinction targeted at most heavily calcified guilds |
| calc_l2h | Primary extinction targeted at least heavily calcified guilds |
| gen_l2h | Primary extinction targeted at specialist guilds |
| gen_h2l | Primary extinction targeted at generalist guilds |
| vuln_l2h | Primary extinction targeted at least heavily predated guilds |
| vuln_h2l | Primary extinction targeted at most heavily predated guilds |

Extinction simulations based on tiering (with extinction selectivity infaunal>pelagic) also produced the closest matches to the empirical post extinction community in terms of community structure and dynamics in 3 out of 13 of the metrics used (i.e. vulnerability, mean trophic level, S1: number of linear chains). Extinction selectivity based on tiering also produced the closest to the post-extinction community in terms of vulnerability when extinction selectivity was reversed (i.e. pelagic>infaunal), but this was only marginally better than extinction selectivity based on tiering (infaunal>pelagic) (Fig 4). Simulations based on extinction selectivity against tiering from infaunal to pelagic also produced relatively close matches to the empirical community in all other metrics apart from generality (Fig. 4).

#### Focus: generality

Scenarios with primary extinction selection based on generality (i.e. the number of resource connections per guild) more closely replicate the post-extinction community than random primary extinction selection with selectivity based both ways (i.e. high to low and low to high) producing almost equally good matches to the empirical data (Fig. 4A). This is likely because extinction selectivity was strongest in the benthic realm, which was made up of taxa that either (i) were primary consumers (i.e. bivalves, brachiopods, crinoids) and only fed upon the basal node of the food web (i.e. low generality) or (ii) were intermediate consumers (i.e. crustaceans, gastropods) that had a broad trophic niche and were feeding upon multiple primary and secondary consumer nodes (i.e. high generality).

Simulations based on selection on generality produced less accurate results than tiering in general, but provided the closest match to the empirical post-extinction community in terms of generality, maximum trophic level (when extinction selectivity is high>low) and diameter, betweeness, S2: omnivory (when extinction selectivity is low>high) (Fig. 4). These results add confidence to the TSS results which show that extinction selectivity was strongest in the benthic realm (where taxa of lowest and highest generality are found) in which the effects of dysoxia/anoxia would be strongest felt.

#### Focus: motility

The only other simulated extinction scenarios that produced close matches to the empirical post-extinction community in terms of structure and dynamics were based on motility (with extinction selectivity non-motile>fast; which produced the closest match for connectance, mean degree, system omnivory index, S4: apparent competition, and S5: direct competition; Fig. 4). This suggests that motility also had a strong bearing on extinction selectivity across the ETE, with non-motile taxa being more highly prone to extinction than motile taxa. Whilst it is evident that the ETE sees the elimination of almost all the motile benthos in the Cleveland Basin, there are also catastrophic losses amongst the non-motile to facultatively motile benthos with the post-extinction community being dominated by 2 to 3 non-motile and facultatively motile taxa with low-oxygen requirements, similarly to modern day low-oxygen ecosystems^38^. However, simulation scenarios of primary extinction selection based on motility perform no better than random primary selection scenarios in regard to guild occupancy in the post-extinction community, thus suggesting that other traits (i.e. tiering) were more important in determining primary extinction vulnerability.

#### Focus other possibilities

Interestingly, some traits that have previously been identified as key determinants of extinction across hyperthermal events, i.e. body size and calcification^14,56^, did not produce simulated post-extinction communities that were a closer match to the empirical post-extinction communities that random selection (Fig. 4). This suggests that the main extinction driver (i.e. dysoxia/anoxia) was not selective based on body size to the same degree as selectivity based on tiering and/or motility. The lack of apparent primary extinction selection based on calcification also suggests that ocean acidification was not a major extinction driver across the ETE in the Cleveland Basin.

### Ecosystem recovery following the ETE

#### Early recovery

The early recovery interval sees an increase in richness as a number of guilds return that were absent from the basin during the post-extinction interval. This return of guilds is associated with the re-oxygenation of the benthic realm (Fig. 2A). However, despite the return of some species occupying motile benthic and infaunal guilds, the majority of new species occupy guilds that were also present during the immediate post-extinction interval (i.e. surficial suspension feeders and pelagic predators). The lower stratigraphic sections from the early recovery interval are characterised by abundant *Dacromya ovum*, a shallow infaunal, low oxygen tolerant, mining bivalve. It has been hypothesised that *D. ovum* may have been able to survive epifaunally before oxygenation improved within the sediment and then subsequently acted as an ecosystem engineer that catalysed the re-oxygenation of the sediment via bioirrigation^38^. *D. ovum* is representative of the first infaunal guild to reappear, some 1 million years after the extinction event^38,39^, and is then followed by subsequent shallow and deep infaunal taxa by the end of the early recovery interval. Whilst the epifaunal community is still dominated by sessile or facultatively mobile taxa, oxygenation of bottom waters is further indicated by the return of motile predatory and grazing guilds (i.e. gastropods and echinoids) as well as the establishment of a greater diversity of soft-bodied epifaunal grazing and infaunal mining guilds (i.e. trace fossils).

Metrics of food web structure suggest that ecosystem recovery is also well underway in the early recovery interval. This period sees connectance and generality returning to lower pre-extinction levels, in-line with increased guild diversity (Fig. 2B and D). Despite a significant recovery of guild diversity and some structural metrics returning towards pre-extinction levels, full ecosystem recovery does not appear to have happened in the early recovery interval. This is evidenced by a paucity of infaunal tiering and motile benthos, as compared to the pre-extinction interval and several of the structural metrics and motifs remain at similar levels to the post-extinction interval rather than starting to return to pre-extinction levels. For example, maximum trophic level and vulnerability (Fig. 2C and E) remain very high and all the food web motifs remain at levels closer to the post-extinction interval than the pre-extinction interval (Fig. 3). This suggests that the early recovery community is still tall, thin and top-heavy with somewhat restricted energy flows consisting of a diverse assemblage of pelagic predators feeding on a still relatively depauperate assemblage of lower-level consumers. This pattern is in contrast to some previous models of ecosystem recovery following mass extinctions that postulate that more-basal trophic levels recovered more quickly than upper trophic levels^10^. Instead, this pattern supports the hypothesis of delayed benthic ecosystem recovery following mass extinctions even in the face of seemingly recovering taxonomic diversity^11^.

#### Late recovery

The late recovery interval witnesses a further increase in guild richness and sees all the structural metrics and motifs return, or start to return, to levels seen in the pre-extinction community (Figs. 2 and 3). Although many of the taxa are different (at species level) to those of pre-extinction community, the majority of pre-extinction guilds are re-occupied by the late recovery interval. Connectance, generality, and vulnerability (Fig. 2B, D-E) are now at levels comparable to the pre-extinction community, as are levels of omnivory (S2) and apparent (S4) and direct (S5) competition (Fig. 3B-D). This suggests that intraguild diversity and functional redundancy is recovering – the reconstructed network indicates a greater number of predators are feeding upon a greater number of prey species thus increasing competition for prey and predator choice simultaneously. The recovery of lower and intermediate-level consumers in the benthic realm drove an increase in the number of linear chains and omnivory, although these metrics are still distinctly lower than the levels seen in the pre-extinction community (Fig. 3A-B).

Together with a further rise in maximum trophic level, these changes in the late recovery phase suggests that the overall shape of the food web still remains much taller and thinner than in the pre-extinction community (Fig. A and Fig 3A). Although the increase in the number of linear chains in the late recovery as compared to the early recovery suggests that food web shape could be starting to return towards pre-extinction levels with the greater diversity of benthic guilds, it may also be a result of changing ecosystem structure brought about by the progression of the Mesozoic Marine Revolution (MMR)^57^. The late recovery interval contains a much more diverse array of benthic predators from groups that were supposedly key drivers of the MMR, such as decapod crustaceans^58^, gastropods^59^ and echinoderms^60^ and such changes in benthic community composition may be driving some of the stepwise increase in maximum trophic level through the system, which deviates from the common pattern of perturbation before return to pre-extinction levels as seen in most of the other structural metrics and motifs (Figs 2 and 3).

### Conclusions

The ETE is characterised by marked changes in community structure and function from a diverse, stable community where each key ecological function is performed by a number of species/guilds to a less diverse, more densely connected and less stable community of generalistic “disaster taxa”^39^ in which nodes (i.e., species/guilds) are more interdependent than in the pre-extinction community. This change from a diverse pre-extinction ecosystem with high degrees of functional redundancy to a contrasting post-extinction community where key functions are performed by single guilds is representative of a “rivet hypothesis”^61^ or a “skeleton crew hypothesis”^8,45^ model in which the subsequent loss of any “rivet” or “crew member” may cause the system to collapse.

Our extinction cascade simulations suggest that primary extinction targeted towards infaunal and epifaunal benthic taxa as well as less motile guilds and extreme generalists or specialists best explain post-extinction community structure and function. These conclusions agree with lithological and geochemical evidence for an anoxia/dysoxia kill mechanism which would primarily target benthic organisms (i.e. sessile suspension feeders or generalist predators) as well as taxa classified as specialists (i.e. those only feeding on a single prey source which is predominantly the basal node of the food web) which are mostly benthic suspension feeders. Despite significant increases in biodiversity during the early recovery interval, most structural and functional metrics suggest ecosystem recovery to pre-extinction levels took at least 7 million years (i.e. until the late recovery interval). However, some metrics suggest that either full recovery had not happened even by the late recovery interval or ecosystem structure and function had re-equilibrated to a different state in the Middle Jurassic and perhaps represents ecological regime shifts associated with the MMR^57,62^.

## Methods

### Dataset

Fossil occurrence data is obtained from a compilation of field data sets^24,26,42,63-65^. The study interval extends from the upper Pliensbachian to the upper Toarcian of the Cleveland Basin (North Yorkshire, UK; Fig 1.) and provides a high resolution data set across the ETE. The data set consists of 38,670 specimens of 162 pelagic and benthic macroinvertebrate species together with occurrences of fish and trace fossils. The data set was subset into four broad time periods; pre-extinction (*margaritatus*-*tenuicostatum* zones of the Staithes Sandstone Formation, Penny Nab and Kettleness Members of the Cleveland Ironstone Formation and Grey Shales Member of the Whitby Mudstone Formation), post-extinction (*serpentinum*-*commune* subzones of the Mulgrave Shale and Alum Shale Members of the Whitby Mudstone Formation), early recovery (upper *bifrons*-lower *levesquei* zones of the Alum Shale, Peak Mudstone and Fox Cliff Siltstone Members of the Whitby Mudstone Formation), and late recovery (upper *levesquei* zone of the Grey and Yellow Sandstone Members of the Blea Wyke Sandstone Formation) (Fig. 1).

### Defining organism ecologies, feeding interactions and trophic guilds

Modes of life were defined for each fossil species based on the ecological traits defined in the Bambach ecospace model^66^ (i.e. motility, tiering, and feeding). Ecological traits were assigned based on interpretations in the published literature which are largely based on functional morphology and information from extant relatives. Information on the body size of each species was also recorded by summarising mean specimen sizes from the section into a categorical classification. The following ecological characteristics were recorded for each fossil species; motility (fast, slow, facultative, non-motile), tiering (pelagic, erect, surficial, semi-infaunal, shallow infaunal, deep infaunal), feeding (predator, suspension feeder, deposit feeder, mining, grazer), and size: gigantic (>500 mm), very large (>300-500mm), large (>100-300mm), medium (>50-100mm), small (>10-50mm), tiny (≤10mm).Size categories are defined by the longest axis of the fossil, estimates of tracemaker size from trace fossils based on literature accounts, or by extrapolating the total length for belemnites from the preserved guard using established approaches^67,68^. A single node for primary producers was added to each food web to ensure that primary consumers were not considered as primary producers in the reconstructions. Feeding interactions were then modelled between organisms based on an inferential model which assigns the possibility of encounter and consumption of prey items using rules defined by inferred ecological foraging traits (i.e., motility, feeding, tiering, and size; Fig. 5). Trophic guilds were defined by unique combinations of ecological and foraging traits (see Supplementary Materials for a full list of trophic guilds and their defining characteristics) which correspond to groups of organisms that have the same predation/prey rules dictating their interactions in the model and are thus akin to trophic species often used in the reconstruction of modern food webs^19,69^. Food webs were produced for each broad time interval (i.e. pre-extinction, post-extinction, early recovery, and late recovery) at both the species and trophic guild level. Further palaeoecological data, which is used to inform the extinction cascade simulations, was also assigned to each trophic species/guild in the food web. This data included motility (fast, slow, facultative, non-motile), tiering (pelagic, epifaunal, infaunal), size (gigantic, very large, large, medium, small, tiny), and calcification (heavy, moderate, light).

**Figure 5.**
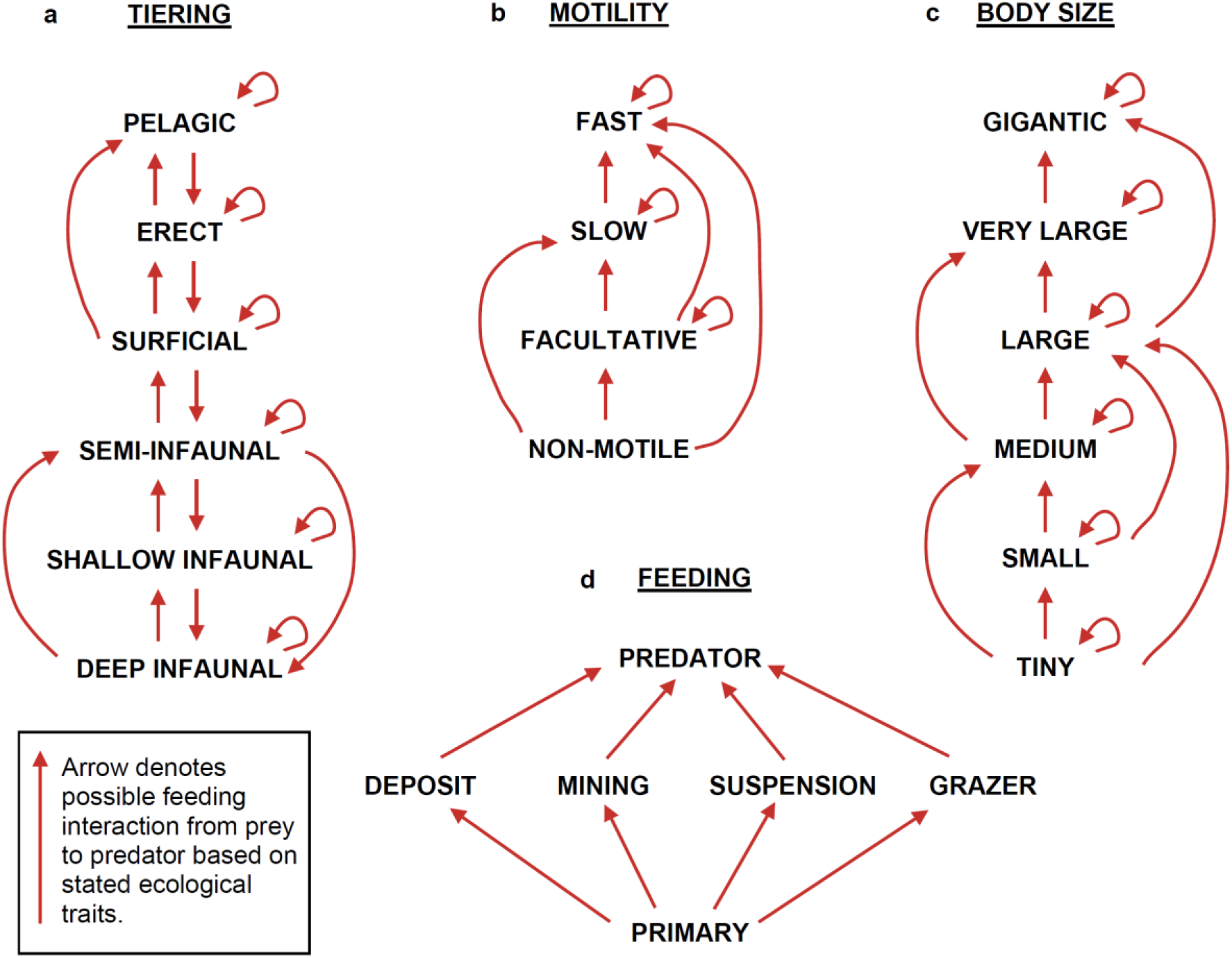
Trait-based feeding rules that parameterise PFIM for reconstructing empirical food webs across all intervals.

### Quantifying community structure and function

Community network structural metrics of size (i.e., richness), connectance (C), maximum trophic level, generality (i.e., in-degree, or number of prey) and vulnerability (i.e., out-degree, or number of predators) as well as the network motifs S1 (i.e. number of linear chains), S2 (i.e. omnivory), S4 (i.e. apparent competition), and S5 (i.e. direct competition) were calculated to track changes in community structure and function across the extinction and through the recovery interval. Food web communities were built, visualised and analysed using the R package *PFIM*.

### Simulating extinction cascades

Extinction cascades were simulated by subjecting guilds in the pre-extinction community to primary extinction scenarios based on ecological and trophic traits that correspond to known vulnerabilities linking the traits to hypothesised mass extinction drivers of anoxia, thermal stress, acidification. For each replicate, we catalogued the timing and identity of all primary extinctions and any secondary extinctions arising when a guild lost all of its resources. The extinction cascades were stopped when the diversity of the simulated post-extinction community reached 21 species and thus equalled that of the empirical post-extinction community.

We explored 13 different scenarios. Simulations were run with primary extinctions selected (i) randomly, (ii/iii) by body size (large to small/small to large), (iv/v) by tiering (infaunal to pelagic/pelagic to infaunal), (vi/vii) by motility (fast to non-motile/non-motile to fast), (viii/ix) calcification (heavy to light/light to heavy), (x/xi) generality (low to high/high to low), and (xii/xiii) vulnerability (low to high/high to low).

We implemented the modelling using the cheddar package in R^70^ using the *RemoveNodes()* function with the ‘cascade’ method of for secondary extinctions. We generated 50 replicates for each scenario by sampling among guilds from within each traits’ levels in the sequence. For example, tiering has six levels (see above) and we randomised the primary extinction sequence of each guild within each of these levels.

Simulated post-extinction food webs were then compared to the empirical post-extinction community using three approaches. First, we compared nine structural metrics between the empirical post-extinction web and the simulated networks. Second, we compared the frequency of four motifs (S1: number of linear chains; S2: number of omnivory motifs; S4: number of apparent competition motifs; S5: number of direct competition motifs) between the empirical post-extinction web and the simulated networks. Third, we used a True Skill Statistic^55^ (TSS/classification-misclassification table/confusion matrix: true positive, true negative, false positive, false negative) to compare the guild-node level similarities of position/identity between the empirical post-extinction web and the simulated networks. All metric calculations were made with functions coded using the *PFIM* package for R.

We combined the inference from all three of these comparisons to identify the most plausible set of primary extinction and associated secondary extinction scenarios (e.g. which trait sequence) that could deliver a community that most closely resembles the post-extinction community.

## Data availability

Contact authors for access to data.

## Code availability

PFIM is currently under publication, contact authors for access to code.

## Acknowledgements

The authors thank the Palaeontological Association for the provision of an Undergraduate Research Bursary (PA-UB01703) to K. Zarzyczny, J. Atkinson, C. Little and A. Dunhill which funded the data collection and initial pilot analysis of this work. We thank the Palaeo@Leeds group for ongoing feedback on this work.

## Author information

A. Dunhill devised the project. K. Zarzyczny, J. Atkinson, C. Little and A. Dunhill collected the data. A. Dunhill, A. Beckerman, J. Shaw and K. Zarzyczny performed the analysis. A. Dunhill lead the write up and all authors contributed to editing and improving the manuscript.

